# Systemic aldehyde storm induced by allyl alcohol exposure results in extensive hepatic ferroptosis in *Aldh2*\**2* knock-in mice

**DOI:** 10.1101/2025.06.18.660242

**Authors:** Yuki Takami, Jun Nakamura, Jun Katahira, Yuhei Maeda, Miyuu Tanaka, Mitsuru Kuwamura, Toshiya Okada, Takeshi Izawa

**Affiliations:** Laboratory of Veterinary Pathology, Osaka Metropolitan University, 1-58 Rinku-Orai-Kita, Izumisano City, Osaka 598-8531, Japan; Laboratory of Laboratory Animal Science, Osaka Metropolitan University, 1-58 Rinku-Ourai-Kita, Izumisano City, Osaka 598-8531, Japan; Laboratory of Cellular Molecular Biology, Osaka Metropolitan University, 1-58 Rinku-Ourai-Kita, Izumisano City, Osaka 598-8531, Japan

**Author notes:** Corresponding author: T. Izawa, Laboratory of Veterinary Pathology, Osaka Metropolitan University, 1-58 Rinku-Orai-Kita, Izumisano City, Osaka 598-8531, Japan.

**Keywords:** acrolein, ALDH2, ferroptosis, glutathione, reactive aldehyde species, rs671

## Abstract

Aldehyde dehydrogenase 2 (ALDH2) is a mitochondrial enzyme that detoxifies multiple aldehyde species in the body. The *ALDH2*2* allele (E487K) is one of the most common gene polymorphisms in humans, resulting in dysfunction of its enzyme activity. This study investigated *in vivo* mechanism of acute liver injury caused by exposure to allyl alcohol (AA) using *Aldh2*2* knock-in (KI) mice with the same amino acid replacement as human *ALDH2*2*. A rapid burst of plasma acrolein as an active metabolite of AA, as well as plasma endogenous reactive aldehydes malondialdehyde (MDA) and formaldehyde, was observed at 10 min after exposure to 75 mg/kg of AA in the *Aldh2*2* KI homozygous mice, which was not evident in the wild-type mice. Hepatocellular necrosis was more extensive in the *Aldh2*2* KI homozygous mice than in the wild-type mice, which was preceded by hepatic glutathione depletion and was coincident with an accumulation of acrolein, MDA, and 4-hydroxy-2-nonenal adducts and iron deposition, suggesting an involvement of ferroptosis in the exacerbation of hepatic necrosis. Recovery from hepatic glutathione depletion was delayed in the homozygous mice compared with the wild-type mice, with a decreasing tendency of hepatic expression of cystine transporter xCT. These results suggest that increased hepatic glutathione consumption, due to decreased aldehyde detoxification capacity, can sensitize the *Aldh2*2* KI homozygous mice to hepatic ferroptosis after the rapid “aldehyde storm”. Our study revealed a crosstalk between aldehyde metabolism and ferroptosis pathways and potential health impacts of endogenous reactive aldehydes on *ALDH2*2* carriers.

## 1. Introduction

Mitochondrial aldehyde dehydrogenase 2 (ALDH2) is a key metabolizing enzyme of acetaldehyde [1]. *ALDH2*2* polymorphism, also known as rs671, is one of the most common enzyme deficiencies in humans, found in 35–45% of East Asians or 560 million of the world population [2–4]. The polymorphism is caused by a single point mutation (G to A nucleotide substitution), leading to a substitution of lysine for glutamate at position 487 (E487K) in the ALDH2 monomer [5]. Heterogenous carriers (*ALDH2*1/*2*) show approximately 40% of ALDH2 enzyme activity of the wild-type (WT) individuals (*ALDH2*1/*1*), while homogenous carriers (*ALDH2*2/*2*) have <1–4% of the WT activity [5–7]. *ALDH2*2* carriers have a high risk for esophageal cancer even with less ethanol consumption [4,6,7] and *ALDH2*2* is also associated with osteoporosis [8], Alzheimer’s disease [9,10], and coronary artery spasm in relation to tobacco smoke exposure [11]. However, the health impacts of *ALDH2*2* on its carriers are still largely unknown.

Reactive aldehyde species, including malondialdehyde (MDA), formaldehyde, and 4-hydroxy-trans-2-nonenal (4-HNE), are derived exogenously from diet and chemicals [12] and also generated endogenously during pathological conditions such as oxidative stress and inflammation [13–17]. The reactive aldehydes form adducts with proteins and DNA, resulting in modification of DNA bases and impaired protein function [13,17–20]. Recent studies have shown that ALDH2 plays an important role in the detoxification of reactive aldehyde species [3,12,19,21]. Therefore, *ALDH2*2* careers with dysfunction of ALDH2 enzyme are expected to be prone to accumulate reactive aldehydes upon exposure. Recent studies have demonstrated that administration of an ALDH2 activator Alda-1 (N-[C1,3-benzodioxol-5-ylmethyl]-2,6-dichlorobenzamide) protects rodents from tissue damage in chronic alcohol consumption-induced atrial fibrillation [22] and hepatic ischemia [23], through a reduction in accumulation of reactive aldehydes. However, effect of Alda-1 on ALDH2 activity is limited to degradation of smaller linear aliphatic aldehydes [24]. Furthermore, *in vitro* studies have shown that Alda-1 also increases expression and activity of ALDHA1, an enzyme involved in the metabolism of 4-HNE and acetaldehyde [25,26]. Therefore, animal models with the gene polymorphism relevant to human *ALDH2*2* are necessary to elucidate the specific mechanism of *ALDH2*2*-related tissue injury.

Acrolein is a highly reactive α,β-unsaturated aldehyde that is generated endogenously during lipid peroxidation [27,28] and also exogenously by metabolic activation of allyl alcohol (AA) in the liver [29,30]. Acrolein can induce hepatotoxicity characterized by periportal (zone 1) necrosis in rodents [30,31]. Acrolein is detoxified by ALDH2 [12,32] or by conjugation with tripeptide thiol glutathione (GSH) in the liver [29,33]. GSH plays a key role in the maintenance of tissue redox homeostasis by scavenging of reactive oxygen species (ROS) and detoxification of lipid peroxides through glutathione peroxidase 4 (GPX4) [16]. However, rapid or mass consumption of GSH stores with acrolein detoxification may lead to decreased antioxidant capacity, which in turn promotes oxidative stress and generation of endogenous aldehydes [14,33–35]. In order to elucidate *in vivo* mechanism of aldehyde-mediated tissue injury in *ALDH2*2* carriers, here we investigate the pathology of *Aldh2*2* knock-in (KI) mice with AA-induced acute exposure to acrolein, focusing on relationship between rapid systemic storm of reactive aldehydes and hepatocellular injury characterized by accumulation of aldehyde adducts.

## 2. Materials and Methods

### 2.1. Animals

The *Aldh2*2* KI mouse strain on C57BL/6N background was generated using CRISPR/CAS9 system by Setsurotech Inc. (Tokushima, Japan). *Aldh2*2* KI mice were obtained by breeding male and female *Aldh2*2* KI homozygous mice while age-matched WT C57BL/6NJcl mice were obtained from CLEA Japan Inc. (Tokyo, Japan). For acute AA toxicity study, 6-week-old male *Aldh2*2* KI homozygous and WT mice were injected intraperitoneally with AA (FUJIFILM Wako Pure Chemical, Osaka, Japan) at the doses of 0 (saline control), 35, 50, and 75 mg/kg and were examined at 24 h post-injection. Mice were euthanized under deep isoflurane anesthesia, and the blood from the caudal vena cava and visceral organs were collected. For toxicokinetic study of AA and acrolein, 6-week-old male *Aldh2*2* KI homozygous and WT mice were injected intraperitoneally with 75 mg/kg of AA and were examined at 0, 10, 30, 60, and 120 min post-injection. Mice were maintained in a room at 21±3°C with a 12 h light-dark cycle and were fed a standard diet (DC-8; CLEA Japan Inc.). Food and water were supplied *ad libitum*. All experimental procedures were approved by the Institutional Animal Care and Use Committee (code nos. 22-154, 23-116, and 24-060) and were performed according to the Guidelines for Animal Experimentation at Osaka Metropolitan University.

### 2.2. Biochemical analyses

ALDH2 enzymatic activity was measured in the liver in order to confirm whether the *Aldh2*2* KI homozygous and heterozygous mice have a decreased ALDH2 activity relevant to human *ALDH2*2* carriers. Isolation of mitochondria from mouse livers was performed as described by Liao *et al* [36]. ALDH2 activity in mitochondrial extracts was measured using XTT (Polyscience Inc, Warrington, PA, USA) with acetaldehyde as substrate according to a previously described assay [37]. Serum samples were subjected to alanine transaminase (ALT) assay with a Transaminase C2 test (Wako Pure Chemical) using a colorimetric method.

### 2.3. Histopathology and immunohistochemistry

The liver, kidneys, heart, lungs, spleen, brain, stomach, intestines, and hind limb including bone, knee joint, and bone marrow, were fixed in 10% neutral buffered formalin for less than 24 h or were fixed in periodate-lysine-paraformaldehyde (PLP) solution [38] at 4°C for 6 h. Fixed tissues were routinely processed, embedded in paraffin, sectioned at a thickness of 3 µm, and were stained with hematoxylin and eosin (HE) for histopathological examination and with Perls’ Prussian blue for detection of iron accumulation. Histopathological evaluation was performed by two board-certified veterinary pathologists (Y.T. and T.I., diplomates JCVP). Paraffin sections with formalin or PLP fixation were used for immunohistochemistry for acrolein (acrolein-lysine adduct), perilipin 2, phosphorylated-histone H2AX (γ-H2AX), and high mobility group box protein 1 (HMGB1) as listed in Table 1. After pretreatment, sections were incubated with 5% skimmed milk for 15 min, followed by 1-h incubation with primary antibodies. After treatment with 3% H_2_O_2_ for 15 min to quench endogenous peroxidase activity, sections were incubated with horseradish peroxidase-conjugated secondary antibody (Histofine simple stain MAX PO^®^; Nichirei Bioscience, Tokyo, Japan) for 1 h and then with 3, 3’-diaminobenzidine (DAB; Nichirei Bioscience) for 10 min. Sections were lightly counterstained with hematoxylin. As a negative control, sections were treated with mouse or rabbit nonimmunized IgG (Agilent Technologies, Santa Clara, CA, USA) instead of the primary antibody. The number of γ-H2AX-positive hepatocytes was counted from randomly selected ten 40× fields (equal to 2.37 mm^2^) of the left lateral lobe in each animal. Formalin-fixed paraffin sections were also subjected to terminal deoxynucleotidyl transferase dUTP nick end labeling (TUNEL) method for evaluation of cell death as previously described [39].

**Table 1.** The list of antibodies used for immunohistochemistry.

| Antibody (clone) | Type | Dilution | Pretreatment | Catalogue No./ Source |
| --- | --- | --- | --- | --- |
| Acrolein (5F6) | Mouse monoclonal | 1:200 | Autoclave in citrate buffer for 10 min | MAR-020n/ JaICA, Shizuoka, Japan |
| Perilipin2 | Rabbit Polyclonal | 1:200 | Autoclave in citrate buffer for 10 min | 15294-1-AP/ Proteintech, Rosemont, IL, USA |
| Phospho-Histone H2A.X (Ser139) | Rabbit monoclonal | 1:500 | Autoclave in citrate buffer for 10 min | 9718/ Cell Signaling Technology, Beverly, MA, USA |
| HMGB1 | Rabbit Polyclonal | 1:200 | Autoclave in citrate buffer for 10 min | ab18256/ Abcam, Cambridge, MA, USA |
| MDA (1F83) | Mouse monoclonal | 1:200 | No pretreatment | MMD-030n/ JaICA |
| 4-HNE (HNEJ-2) | Mouse monoclonal | 1:200 | No pretreatment | MHN-020P/ JaICA |

### 2.4. Specific *in situ* detection of aldehyde adducts on frozen tissue sections

Formalin-based solutions are most commonly used as a fixative for immunohistochemistry for diagnostic and research purposes; however, such fixatives may form artificial formaldehyde adducts on tissue specimens as formaldehyde can react with arginine, cysteine, tryptophan, lysine, histidine, and the *N*-terminal amine to form methylol adducts [40–42]. For specific *in situ* detection of aldehyde adducts, ten-μm fresh-frozen sections of the left lateral lobe of the liver were cut on a cryostat and fixed in formalin-free zinc chloride fixative (Catalogue #550523, BD Biosciences Pharmingen, Heidelberg, Germany) at room temperature for 15 min. The sections were then immunostained with anti-MDA (clone 1F83) or anti-4-HNE (clone HNEJ-2) antibody (Table 1) as in immunohistochemistry using paraffin sections described above.

### 2.5. High performance liquid chromatography (HPLC) analysis for plasma aldehydes measurement

All HPLC analyses were conducted using an Agilent 1260 Series chromatography system (Agilent Technologies) equipped with both a photodiode array detector and a fluorescence detector. Chromatographic separation was carried out on a COSMOCORE C18 column (4.6 mm i.d. × 50 mm, 2.6 μm; Nacalai Tesque, Kyoto, Japan). Commercially available methanol-free formaldehyde (Thermo Fisher Scientific Inc., Waltham, MA, USA), malondialdehyde bis(dimethyl acetal) (Sigma-Aldrich, Merck KGaA, Darmstadt, Germany), and acrolein (Tokyo Chemical Industry Co., Ltd., Tokyo, Japan) were used as standards. These compounds were serially diluted with water and then derivatized to prepare calibration curves. Plasma samples were first deproteinized with perchloric acid, followed by derivatization and HPLC analysis as previously described. Formaldehyde was derivatized with dimedone to form formaldehyde-meton and detected using HPLC with a photodiode array detector [43]. MDA and acrolein were derivatized with 2-thiobarbituric acid [44] and 3-aminophenol [45], respectively, and analyzed by HPLC with a fluorescence detector.

### 2.6. Western blot

Liver samples from the left medial lobe were homogenized in a RIPA buffer with proteinase inhibitor cocktail (Nacalai tesque). After centrifugation at 13,000 × g for 10 min, the supernatant was mixed with an equal volume of 2× SDS sample buffer (125 mM Tris-HCl, pH 6.8, 4% SDS, 30% glycerol, and 10% 2-mercaptoethanol) and then boiled at 95°C for 5 min. The protein concentration was determined by an absorption spectrometer using Bio-Rad Protein Assay (Bio-Rad Laboratories, Hercules, CA, USA). Samples were separated on 5-20% gradient polyacrylamide gels (ATTO Corporation, Tokyo, Japan) and transferred to polyvinylidene difluoride membranes (Bio-Rad Laboratories). The membranes were incubated overnight at 4°C with primary antibodies against ALDH2, alcohol dehydrogenase 5 (ADH5), GPX4, glutamate-cysteine ligase catalytic subunit (GCLC), cystine transporter solute carrier family 7 member 11/xCT (xCT), ferritin, and transferrin receptor 1 (TFR1) as listed in Table 2, followed by an incubation with peroxidase-conjugated secondary antibody (Histofine Simple Stain MAX PO^®^; Nichirei Bioscience) for 30 min. Signals were visualized with ECL Prime (Cytiva, Marlborough, MA, USA), and scanned with a luminescent image analyzer (LAS-4000; GE Healthcare, Madison, WI, USA). The data were normalized by total protein bands on gels stained with Coomassie brilliant blue (Fig. S1) as described previously [46].

**Table 2.**
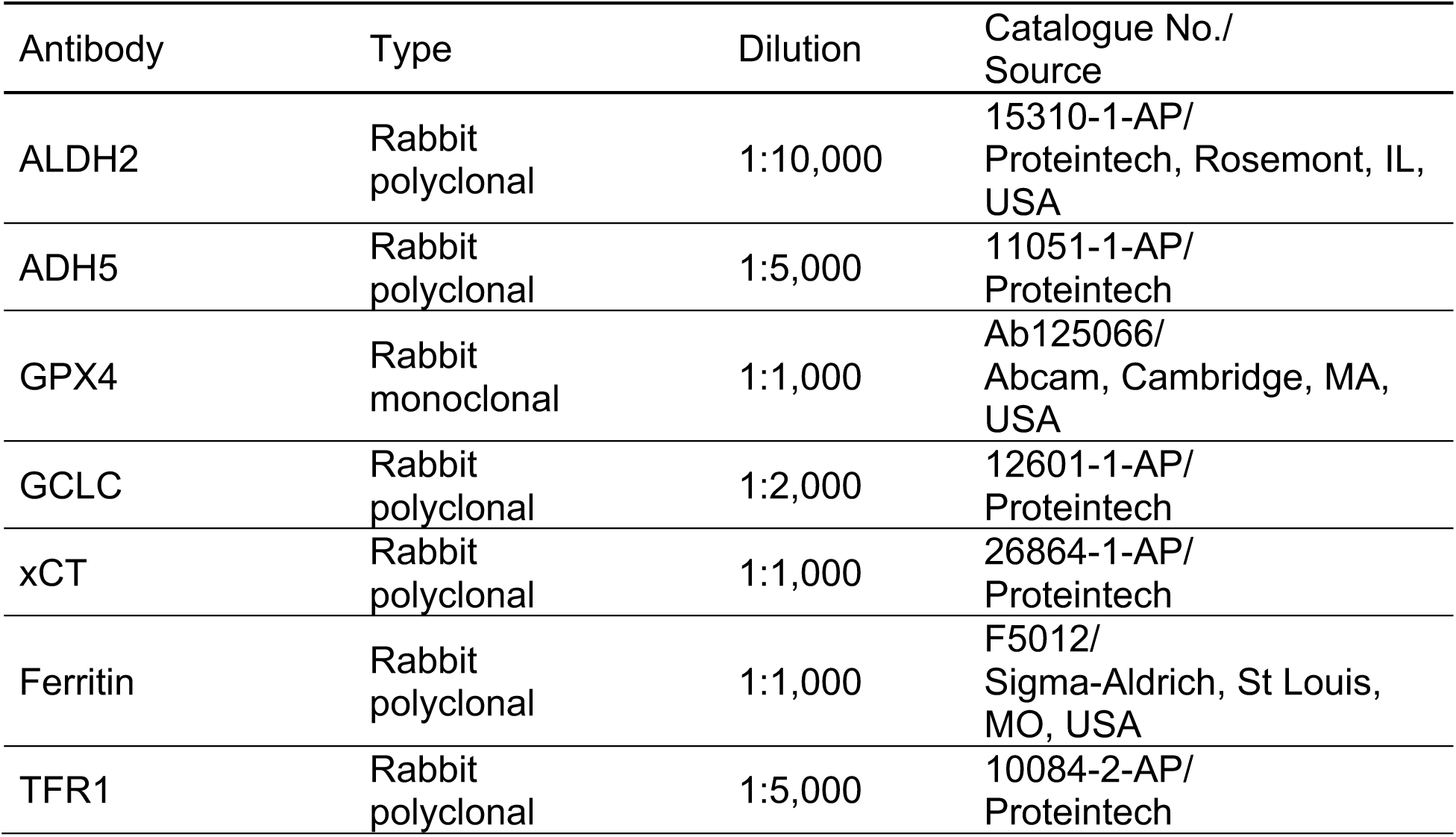
The list of antibodies used for Western blot.

### 2.7. Quantification of hepatic GSH and MDA

Hepatic content of GSH and glutathione disulfide (GSSG) was measured using the GSSG/GSH Quantification kit (Dojindo, Kumamoto, Japan) in accordance with the manufacturer’s instructions. The GSSG/GSH ratio is regarded as an indicator of tissue redox status [47]. Hepatic MDA content was measured by thiobarbituric acid reactive substances (TBARS) method using an MDA assay kit (Northernwest Life Science Specialities, Vancouber, Canada) according to the manufacturer’s instructions.

### 2.8. Statistical analysis

Data obtained were expressed as mean ± standard deviation (SD). Statistical analyses were performed using two-way analysis of variance (ANOVA) followed by Tukey’s multiple comparison by a Prism software (ver. 10.3.0.; GraphPad, San Diego, CA, USA). Significance was considered at *P* < 0.05.

## 3. Results

### 3.1. The *Aldh2*2* KI mice have a decreased ALDH2 enzyme activity relevant to human *ALDH2*2* carriers

The *Aldh2*2* KI mouse, which has an *Aldh2* gene mutation orthologous to the human *ALDH2*2*, was generated by genome editing of 1516G in the exon 12 of the endogenous murine *Aldh2* locus, via inserting a G-to-A substitution (E504K) by CRISPR-Cas9 system (Fig. 1A). The KI allele was detected by sequencing of genomic PCR products including the G1516A site (Fig. 1B). The enzymatic activity of ALDH2 was significantly decreased in the liver of *Aldh2*2* KI homozygous and heterozygous mice compared with that of WT mice, which is relevant to human *ALDH2*2* carriers (Fig. 1C). Anatomical appearance (Fig. 1D) and body weight (Fig. 1E) did not differ between the *Aldh2*2* KI homozygous and WT mice.

**Figure 1.**
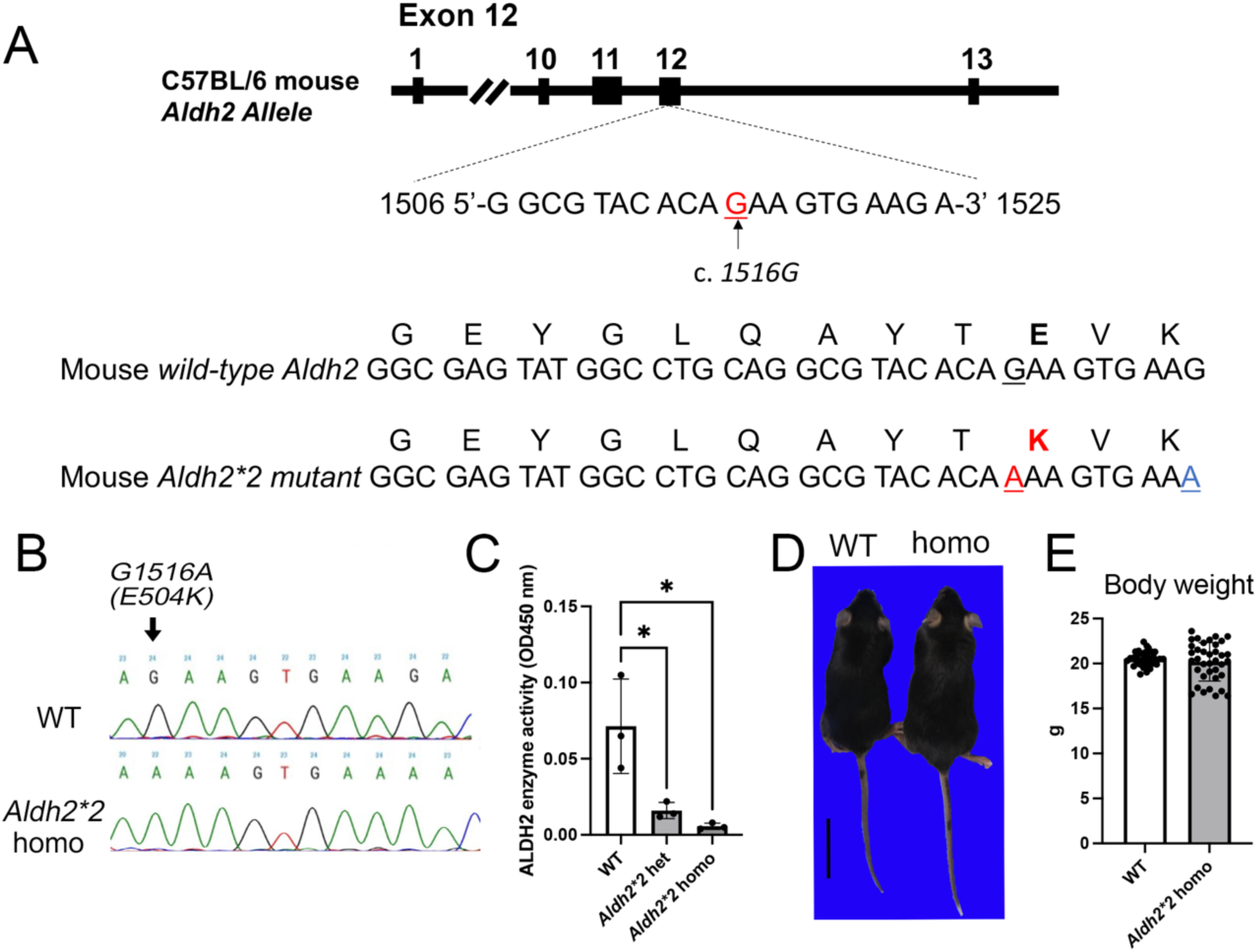
Generation of *Aldehyde dehydrogenase 2*2* (*Aldh2*2)* knock-in mouse. (A) The target site of 1516G is located in the exon 12 of the mouse *Aldh2* gene. A G-to-A substitution at this site in the mouse *Aldh2*2* allele, resulting in a Glu504Lys substitution (highlighted in red), is homologous to the point mutation (rs671) in the human *ALDH2*2* allele. Silent mutation of the G1524A (highlighted in blue) is designated to confirm successful knock-in (KI) of the target sequence. (B) Genotyping of the *Aldh2*2* KI mice were performed with genomic PCR and direct sequencing that can detect the G1516A mutation of the mouse *Aldh2* gene. (C) Hepatic ALDH2 enzyme activity is markedly decreased in the *Aldh2*2* KI homozygous and heterozygous mice compared with wild-type (WT) mice. * *P*<0.05 vs. WT mice, by Tukey’s multiple comparison (n=3 in each genotype). (D) Gross appearance of the body is similar between the WT and *Aldh2*2* KI homozygous mouse. Bar=3 cm. (E) Body weight does not differ significantly between the male WT and homozygous mice (n=33 in the WT and n=37 in the homozygous mice) at the age of 6 weeks.

### 3.2. AA exposure induces acute and systemic bursts of reactive aldehydes in the *Aldh2*2* KI mice

AA is metabolized in the liver to highly-reactive acrolein, which is further detoxified by ALDH2 and conjugation to GSH with or without catalyst [12,29,48]. Plasma acrolein levels peaked at 10 min after AA exposure at 75 mg/kg, with the peak levels being much higher at 10 and 30 min in the *Aldh2*2* KI homozygous mice than in the WT mice (Fig. 2A), suggesting a decreased capacity of acrolein detoxification in the *Aldh2*2* KI mice. Plasma levels of endogenous aldehydes MDA and formaldehyde did not change significantly after AA exposure in the WT mice (Fig. 2B and C). In contrast, plasma MDA levels increased at 10 to 60 min after AA exposure in the *Aldh2*2* KI homozygous mice (Fig. 2B). Plasma formaldehyde levels were also significantly higher at 10 to 60 min in the homozygous mice than in the WT mice (Fig. 2C) and interestingly, they remained high at 120 min in the homozygous mice. These results suggest that AA exposure to the *Aldh2*2* KI mice induces a rapid and systemic storm of reactive aldehydes that includes not only the AA metabolite acrolein but also endogenous reactive aldehydes MDA and formaldehyde.

**Figure 2.**
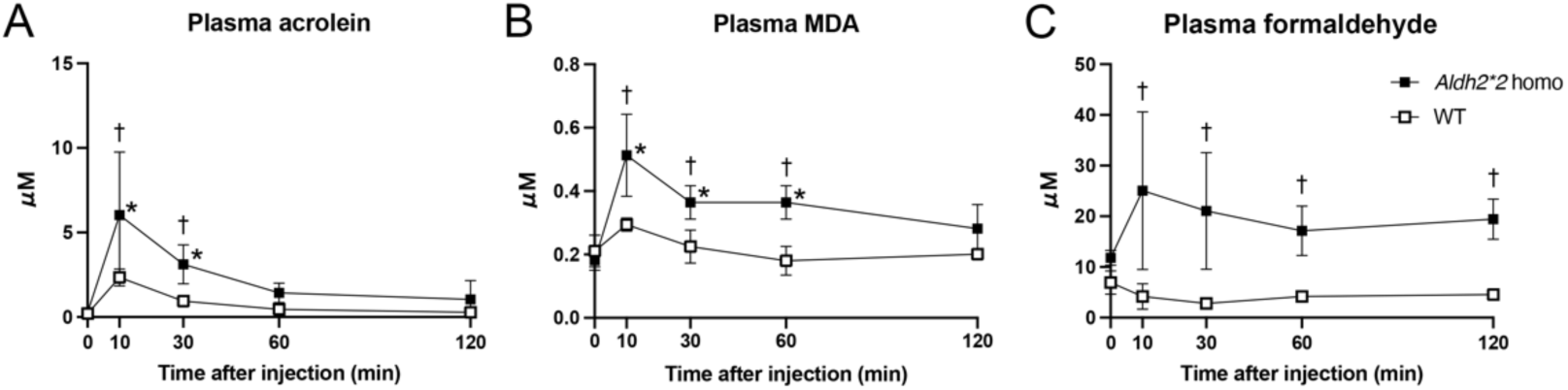
Allyl alcohol (AA) administration induces a rapid and systemic aldehyde storm in the *Aldh2*2* KI homozygous mice. Kinetics of plasma (A) acrolein, (B) malondialdehyde (MDA), and (C) formaldehyde after AA exposure in the WT and *Aldh2*2* KI homozygous mice (n=4 in each time point). Plasma levels of acrolein, MDA, and formaldehyde increases from 10 min after AA exposure in the *Aldh2*2* homozygous mice compared with WT mice. * *P*<0.05 vs. 0 min (treatment factor); † *P*<0.05 vs. WT mice of the same time point (*Aldh2*2* factor), by Tukey’s multiple comparison.

### 3.3. Acute liver injury after AA exposure is deteriorated in the *Aldh2*2* KI mouse with an increased accumulation of aldehyde adducts

Pathological examination revealed that the systemic storm of reactive aldehydes, induced by AA exposure, affected the spleen, bone marrow, small intestine, and liver of the *Aldh2*2* KI homozygous mice. The liver is most severely affected among them and histopathological findings of non-hepatic organs include decreased density of hematopoietic cells with increased nuclear debris in the bone marrow, increased apoptosis of the splenic lymphoid follicle, and degeneration/single cell necrosis of crypt epithelium of the small intestine (Figs. S2A and B). Immunohistochemistry for γ-H2AX, a specific marker for DNA double strand breaks [49], revealed increased nuclear immunoreactivity in these affected cells (Fig. S2C), indicative of DNA damage.

Serum ALT levels significantly increased in the *Aldh2*2* KI homozygous mice at 24 h after exposure to 75 mg/kg AA, compared with those in the WT mice (Fig. 3A). Grossly, focal or spotty, pale discoloration of the liver was observed in the WT mice at 75 mg/kg (Fig. 3B) and the *Aldh2*2* KI homozygous mice at 50 mg/kg. The pale lesions expanded throughout the whole liver in the *Aldh2*2* KI homozygous mice at 75 mg/kg (Fig. 3B). Histopathologically, periportal to massive coagulative necrosis with vacuolar degeneration of hepatocytes was seen in the liver of the WT mice at 75 mg/kg (3 of 7 mice) and the *Aldh2*2* KI homozygous mice at 50 mg/kg (1 of 3 mice) and 75 mg/kg (11 of 11 mice) at 24 h after AA exposure (Table 3 and Fig. 3C). Earliest histopathological lesions, which appeared as a vacuolar degeneration of hepatocytes, were detected at 30 min in the liver of the homozygous mice (Fig. S3). Such degeneration was also present in the WT mice from 60 min onward. Hepatocyte necrosis was seen at 120 min in a half of the *Aldh2*2* KI homozygous mice while necrosis was absent in the WT mice until 120 min.

**Figure 3.**
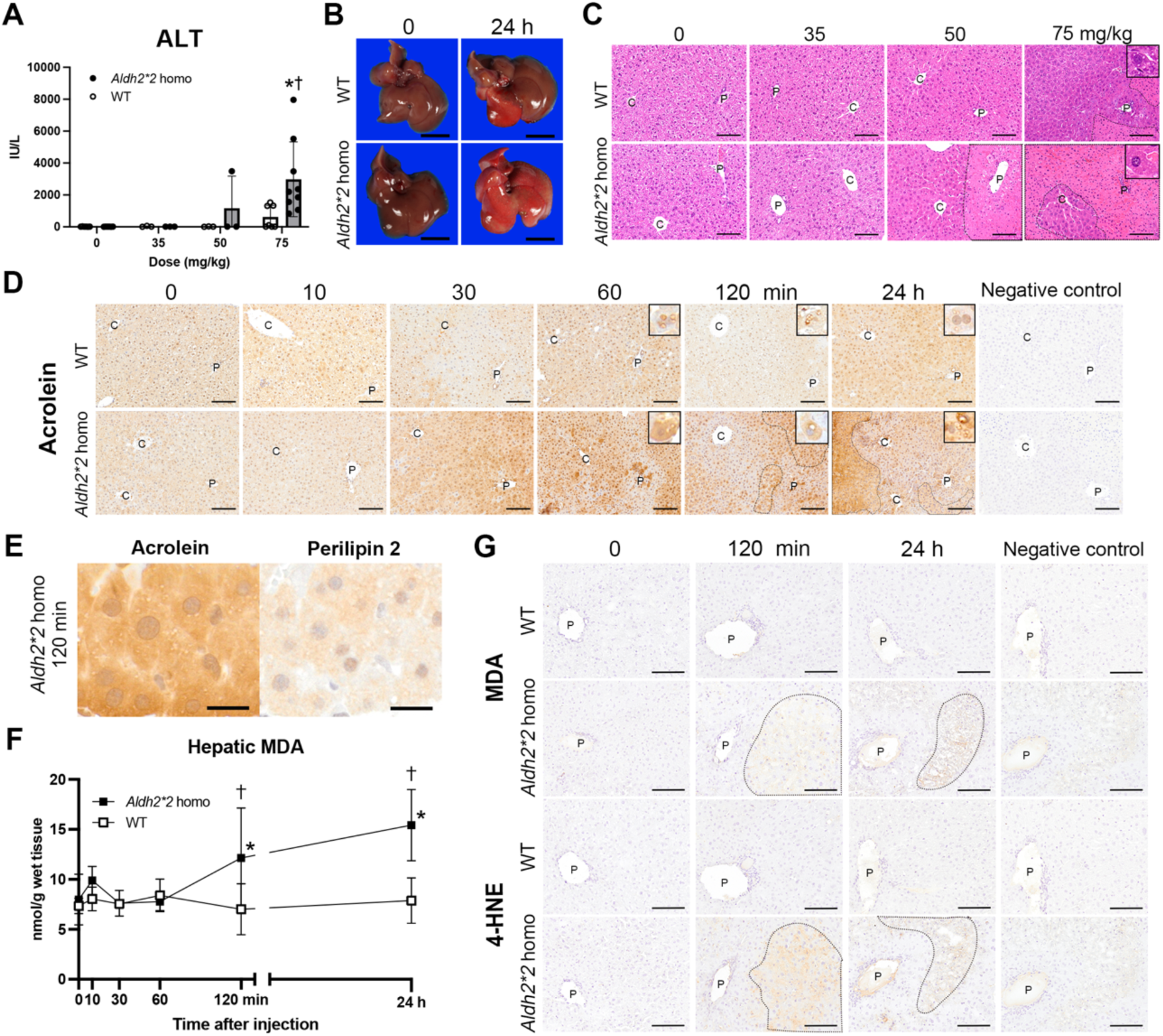
Acute liver injury after AA exposure is markedly exacerbated with an accumulation of reactive aldehydes in the *Aldh2*2* KI homozygous mice. (A) Serum alanine aminotransferase (ALT) levels in the WT and *Aldh2*2* KI homozygous mice at 24 h after AA administration at the doses of 0 (saline), 35, 50 and 75 mg/kg. Serum ALT levels are significantly higher in the homozygous mice than in the WT mice at 75 mg/kg. * *P*<0.05 vs. 0 mg/kg group (treatment factor); † *P*<0.05 vs. WT mice of the same dose (*Aldh2*2* factor), by Tukey’s multiple comparison. (B) Gross appearance of the liver of the mice injected with 0 and 75 mg/kg of AA. The liver has focal or spotty, pale discoloration at 24 h after AA exposure, with the extent being greater in the *Aldh2*2* KI homozygous mice than in the WT mice. Bar=2 cm. (C) Histopathology of the liver at 24 h after AA exposure. Periportal to massive coagulative necrosis (encircled by dotted line) with vacuolar degeneration of hepatocytes (insets) are observed in the WT mice at 75 mg/kg and in the *Aldh2*2* KI homozygous mice at 50 and 75 mg/kg. The extent of hepatic necrosis is much greater in the homozygous than in the WT mice at these doses. Hematoxylin and eosin stain. Bar=100 µm. (D) Immunohistochemistry for acrolein adducts in the liver during development of acute liver injury. Necrosis area, encircled by dotted line, is intensely positive for acrolein adducts. Note that the cytoplasmic vacuoles in the degenerative hepatocytes are often immunolabeled for acrolein adducts (insets). Bar=50 µm. C; central vein, P; portal vein. (E) Immunohistochemistry for acrolein adducts and perilipin 2 using serial sections of the liver of the *Aldh2*2* KI homozygous mice at 120 min after AA exposure at 75 mg/kg. Bar=20 µm. (F) Hepatic MDA contents measured with the thiobarbituric acid reactive substances (TBARS) method. Hepatic MDA significantly increases in the homozygous mice from 120 min after AA exposure (n=3 in the WT at 30 min and n=4 in other time points) * *P*<0.05 vs. 0 min (treatment factor); † *P*<0.05 vs. WT mice of the same time point (*Aldh2*2* factor), by Tukey’s multiple comparison. (G) Immunohistochemistry for MDA adducts (clone 1F83) and 4-hydroxy-2-nonenal (4-HNE) adducts (clone HNEJ-2) shows an accumulation of these adducts in the necrotic/degenerative lesions (encircled by dotted line) of the liver of the homozygous mice. Bar=100 µm.

**Table 3.** Histologic scoring for acute liver injury after allyl alcohol exposure.

| Degeneration/necrosis, hepatocyte, mainly zones1-2 |  |  |  |  |  |  |  |  |
| --- | --- | --- | --- | --- | --- | --- | --- | --- |
| Score | saline |  | 35 |  | 50 |  | 75 mg/kg AA |  |
|  | WT | homo | WT | homo | WT | homo | WT | homo |
| 0 (absent) | 7 | 6 | 3 | 3 | 3 | 2 | 4 | 0 |
| +1 (mild) | 0 | 0 | 0 | 0 | 0 | 0 | 0 | 0 |
| +2 (moderate) | 0 | 0 | 0 | 0 | 0 | 0 | 1 | 5 |
| +3 (severe) | 0 | 0 | 0 | 0 | 0 | 1 | 2 | 6 |

Histopathological score 0 (absent), normal histological hepatic architecture without histological abnormality; +1 (mild), focal degeneration of hepatocytes; +2 (moderate), extensive degeneration of hepatocytes with occasional necrosis or focal coagulative necrosis of hepatocytes; +3 (severe), massive or extensive coagulative necrosis and degeneration of hepatocytes. The value represents the number of individuals classified by the score.

Immunohistochemistry with anti-acrolein antibody (clone 5F6) was performed for detection of acrolein bound to lysine residues of proteins (acrolein-lysine adducts) within the hepatic lesions (Fig. 3D). At 10 min after exposure to AA at 75 mg/kg, individual hepatocytes with accumulation of acrolein adducts were scattered in the liver of the WT and *Aldh2*2* KI homozygous mice. The immunoreactivity for acrolein adducts was markedly increased in the *Aldh2*2* KI homozygous mice at 60, 120 min, and 24 h compared with the WT mice, with an accumulation of the acrolein adducts in the necrotic areas. Additionally, the cytoplasmic vacuoles of hepatocytes observed in the HE slides (Fig. S3 and Fig. 3C), were immunopositive for acrolein adducts (Fig. 3D insets). Immunohistochemistry using serial sections revealed that the vacuolar hepatocytes with accumulation of acrolein adducts were immunopositive for perilipin 2 (Fig. 3E), a lipid droplet coat protein [50,51]. These results suggest that the accumulation of acrolein adducts is associated with fatty degeneration in the hepatocytes after acute acrolein exposure.

Hepatic MDA content increased at 120 min and 24 h in the *Aldh2*2* KI homozygous mice compared with the WT mice (Fig. 3F). Consistent with this change, MDA and 4-HNE adducts were accumulated within the degenerative/necrotic hepatocytes at 120 min and 24 h in the *Aldh2*2* KI homozygous mice, but not in the WT mice (Fig. 3G). These results suggest that acute liver injury after AA exposure is deteriorated in the *Aldh2*2* KI homozygous mouse in association with the accumulation of aldehyde adducts following the systemic aldehyde storm.

### 3.4. The acute aldehyde storm increases hepatic GSH consumption in the *Aldh2*2* KI mice with decreased aldehyde detoxification, resulting in increased hepatic ferroptosis

To investigate the cause of the systemic and hepatic aldehyde accumulation in the *Aldh2*2* KI homozygous mice, we investigated hepatic expression of aldehyde detoxification enzymes. Expression levels of hepatic ALDH2 did not change significantly after AA exposure and were much lower in the *Aldh2*2* KI homozygous mice than in the WT mice (Fig. 4A and B). Hepatic expression of ADH5, an enzyme responsible for intracellular formaldehyde detoxification [19,52], were lower after AA exposure in the *Aldh2*2* KI homozygous mice than in the WT mice, despite the persistent increase in the plasma formaldehyde (Fig. 4A and C). We next investigated hepatic GSH metabolism as GSH plays an important role in acrolein detoxification [29,33] as well as regulation of tissue redox homeostasis [16]. Hepatic GSH content rapidly decreased at 10 min after AA exposure with the lowest peak at 30 min in the WT and *Aldh2*2* KI homozygous mice (Fig. 4G). Recovery of the hepatic GSH was delayed in the *Aldh2*2* KI homozygous mice compared with the WT mice; the GSH content tended to be lower in the homozygous than in the WT mice at 24 h (p=0.09). The hepatic GSH content was negatively correlated with the plasma acrolein (r=−0.45, p=0.003) and plasma MDA (r=−0.47, p=0.002). The ratio of oxidized to reduced glutathione (GSSG/GSH ratio) is regarded as an indicator for tissue redox conditions [47,53] and it is increased at 10 to 60 min in the *Aldh2*2* KI homozygous mice compared with the WT mice (Fig. 4H), indicating the induction of oxidative stress in the homozygous liver at the period of the aldehyde storm. The GSSG/GSH ratio was positively correlated with the plasma acrolein (r=0.78, p<0.001), plasma MDA (r=0.73, p<0.001), and plasma formaldehyde (r=0.47, p=0.002), suggesting that the hepatic oxidative stress is closely associated with the systemic aldehyde storm. xCT is a component of system x_c_^−^ cystine/glutamate antiporter that is responsible for the uptake of cystine for intracellular GSH synthesis. Hepatic expression of xCT tended to be lower at 60 min (p=0.080) and was significantly lower at 24 h after AA exposure in the *Aldh2*2* KI homozygous mice than in the WT mice (Fig. 4D). GPX4 is an antioxidant enzyme that detoxifies lipid hydroperoxides using GSH as a co-factor [16,54] and GCLC is a subunit of heterodimeric GSH biosynthetic enzyme [55]. Hepatic expression of GCLC or GPX4 did not change significantly after AA exposure in the WT or *Aldh2*2* KI homozygous mice (Fig. 4A, E and F).

**Figure 4.**
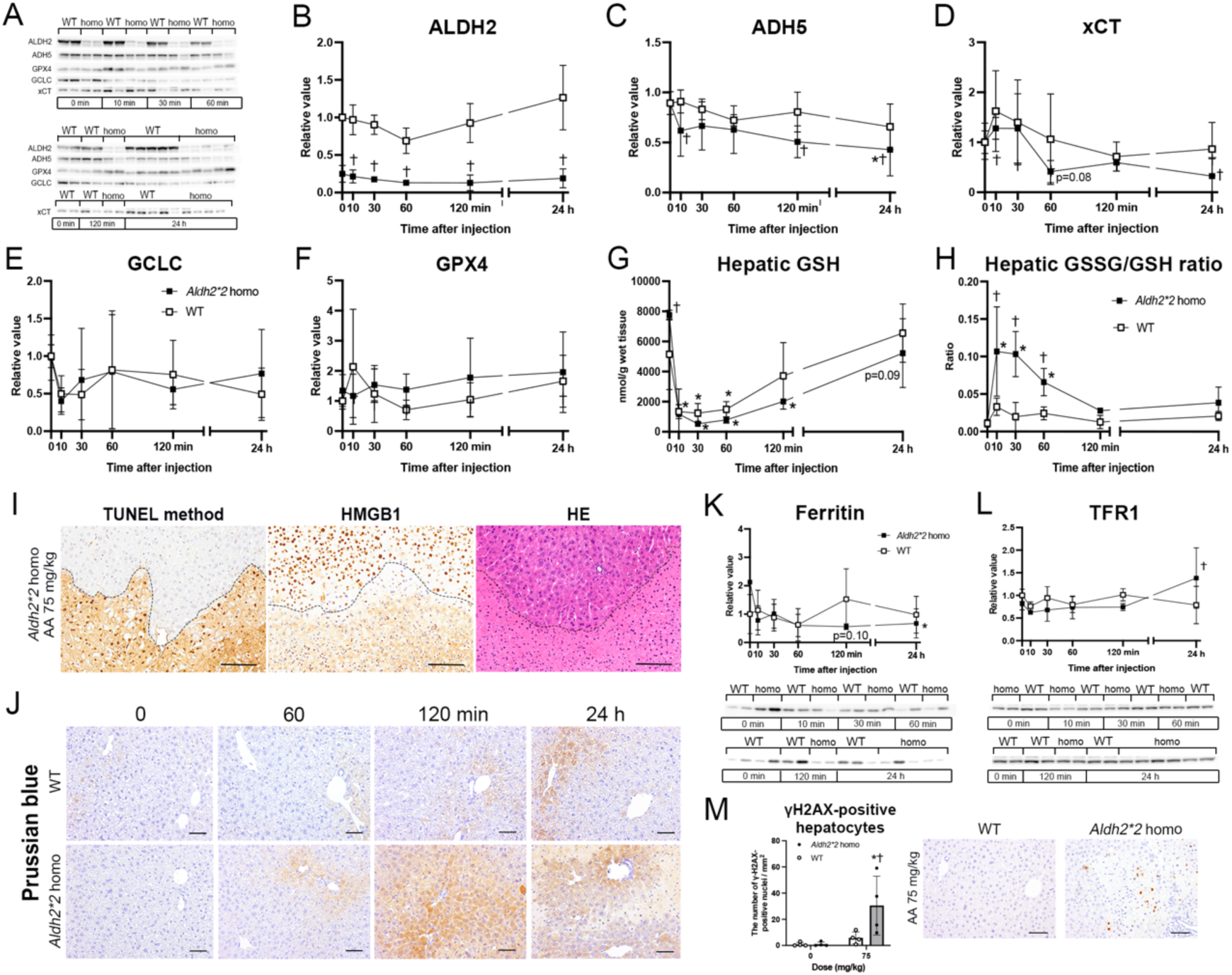
Decreased capacity of aldehyde detoxification increases hepatic glutathione (GSH) consumption, leading to extensive hepatic ferroptosis in the *Aldh2*2* KI homozygous mice. (A-F) Western blot data for (B) ALDH2, (C) aldehyde dehydrogenase 5 (ADH5), (D) cystine transporter solute carrier family 7 member 11/xCT (xCT), (E) glutamate-cysteine ligase catalytic subunit (GCLC), and (F) glutathione peroxidase 4 (GPX4). Hepatic ALDH2 expression is consistently lower in the *Aldh2\**2 KI homozygous mice than in the WT mice, while hepatic ADH5 expression decreases after AA exposure in the homozygous mice (n=7 in the WT and n=11 in the homozygous mice at 24 h; n=4 in other time points). (G) Hepatic GSH content and (H) the ratio of glutathione disulfide (GSSG) to GSH during development of acute liver injury. Hepatic GSH decreases immediately after AA exposure in both the WT and homozygous mice while the recovery from the decreased GSH is delayed in the *Aldh2\**2 KI homozygous mice compared with the WT mice. Hepatic GSSG/GSH ratio increases at 10 to 60 min after AA exposure in the homozygous mice (n=7 in the WT and n=11 in the homozygous mice at 24 h; n=4 in other time points). (I) Representative images of TUNEL method, HMGB1 immunohistochemistry, and HE in the hepatic necrosis lesions of the *Aldh2*2* KI homozygous mice at 24 h after exposure to 75 mg/kg AA. Necrotized hepatocytes (below the dotted line) have an intense nuclear and cytoplasmic labeling for TUNEL and a loss of nuclear high mobility group box protein 1 (HMGB1) immunolabeling with the absent to faint cytoplasmic immunolabeling. (J) Representative images of diaminobenzidine (DAB)-enhanced Perls’ Prussian blue stain in the liver of the WT and *Aldh2*2* KI homozygous mice at 0, 60, 120 min and 24 h after exposure to 75 mg/kg AA. Accumulation of iron is marked in the degenerative/necrotic hepatocytes of the homozygous mice. (K-L) Western blot data for (K) ferritin and (L) transferrin receptor 1 (TFR1) in the liver of the WT and *Aldh2*2* KI homozygous mice (n=7 in WT and n=11 in homozygous mice at 24 h; n=4 in other time points). Hepatic ferritin expression tended to be lower in the homozygous mice than in the WT mice at 120 min after AA exposure. Hepatic TFR1 expression is higher in the homozygous mice than in the WT mice at 24 h after AA exposure. (M) The number of γH2AX-positive hepatocyte nuclei of the WT and homozygous mice 24 h after AA exposure at 0 and 75 mg/kg (n=4 in each group). The number of γH2AX-positive hepatocytes is greater in the homozygous mice at 75 mg/kg than in the WT mice. * *P*<0.05 vs. 0 min or 0 mg/kg group (treatment factor); † *P*<0.05 vs. WT mice of the same time point or dose (*Aldh2*2* factor), by Tukey’s multiple comparison. Bar=50 µm.

Accumulation of lipid peroxides under oxidative stress condition may induce ferroptosis, an iron-dependent regulated cell death [56]. Thus, we investigated the characteristics of the hepatocellular injury in our *Aldh2*2* mouse model. TUNEL method revealed a marked accumulation of DNA fragments in both the cytoplasm and nucleus of necrotic hepatocytes in the *Aldh2*2* KI homozygous mice after AA exposure (Fig. 4I). Cytoplasmic leakage of the DNA fragments is considered to result from nuclear envelope rupture due to activation of proteinases and lipases in necrotic cells [57,58]. HMGB1 is a nonhistone chromosomal protein that is translocated from the nucleus to the cytoplasm and finally released from cells undergoing necrosis [59,60] and mediates inflammation as one of damage-associated molecular patterns (DAMPs) [61,62]. Immunohistochemistry for HMGB1 showed translocation of HMGB1 from the nucleus to the cytoplasm in the necrotic hepatocytes of the *Aldh2*2* KI homozygous mice (Fig. 4I). Next, we evaluated cellular iron deposition in the hepatic lesions as iron accumulation can promote the Fenton reaction that propagates lipid peroxidation [63,64]. Intense iron accumulation was present in the degenerative and necrotic hepatocytes at an early stage of the acute liver injury (120 min after AA exposure) in the *Aldh2*2* KI homozygous mice, while iron accumulation was mild in the WT mice at this time point (Fig. 4J). Ferritin is an intracellular iron storage protein responsible for sequestering free iron and has a protective role against ferroptosis by preventing Fenton reaction [56]. Degradation of ferritin by selective autophagy (ferritinophagy) can sensitize cells to ferroptosis [65]. Hepatic expression of ferritin tended to be lower (p=0.10) in the *Aldh2*2* KI homozygous mice at 120 min after AA exposure than in the WT mice (Fig. 4K). TFR1 is a receptor for transferrin-bound iron and is responsible for cellular iron uptake [66]. Hepatic expression of TRF1 did not change between the WT and *Aldh2*2* KI homozygous mice until 120 min after AA exposure and became greater in the homozygous mice than in the WT mice at 24 h (Fig. 4L). As reactive aldehydes are known to form DNA adducts and to induce DNA damage [67,68], we performed immunohistochemistry for γ-H2AX on the liver sections. The number of γ-H2AX positive nuclei was significantly increased in the hepatic lesions of the *Aldh2*2* KI homozygous mice at 24 h, compared with those in the WT mice (Fig. 4M). These results suggest that the acute and systemic aldehyde storm in the *Aldh2*2* KI mice results in hepatic ferroptosis characterized by extensive necrosis, accumulation of lipid peroxidation-derived aldehydes and iron, and DNA damage. Hepatic GSH depletion, caused by decreased capacity of aldehyde detoxification in the *Aldh2*2* KI mice, is considered to contribute to the extensive hepatic ferroptosis.

### 3.5. Comparison of basal hepatic function between the WT and *Aldh2*2* KI homozygous mice

To characterize physiological hepatic function of the *Aldh2*2* KI mice, we performed comparison of the hepatic parameters between the WT and *Aldh2*2* KI homozygous mice (Fig. S4). The characteristic differences are as follows: The *Aldh2*2* KI homozygous mice have a decreased hepatic ALDH2 expression with a decreasing tendency of hepatic ADH5 expression, increased plasma formaldehyde, increased hepatic MDA, and increased hepatic GSH content compared with the WT mice. These results suggest that the *Aldh2*2* KI mice have an alteration in the aldehyde and glutathione metabolism without any treatment.

## 4. Discussion

The *Aldh2*2* KI mouse strain generated in this study has a single nucleotide substitution that is homologous to the rs671 substitution of human *ALDH2*2* allele. Compared with the *Aldh2* knockout mouse strain with a complete lack of ALDH2 function [3], the *Aldh2*2* KI strain has some residual ALDH2 enzyme activity, which is relevant to the human *ALDH2*2*. Therefore, our *Aldh2*2* KI strain would be a better model to investigate pathological conditions of the human *ALDH2*2* carriers. Recent studies using other *Aldh2*2* KI mouse strains have shown increased ROS production in response to cisplatin treatment [69], greater susceptibility to alcohol-induced atrial fibrillation [22], and increased cardiovascular oxidative stress after aldehyde exposure [70].

Our results showed that acute acrolein exposure, induced by AA administration, is accompanied with systemic burst of endogenous reactive aldehydes including MDA and formaldehyde in the *Aldh2*2* KI mice. The possible mechanism of this “systemic aldehyde storm” is as follows: AA is mainly metabolized in the liver [30], and its metabolite acrolein is detoxified mainly by GSH conjugation catalyzed by glutathione S -transferase or without catalyst and partly by aldehyde dehydrogenases such as ALDH2 [48]. Moreover, acrolein itself can inactivate ALDH2 activity by forming adduct at its NAD binding site [32]. ALDH2 dysfunction in the *Aldh2*2* KI mice leads to decreased capacity of acrolein detoxification and thereby increased consumption of hepatic GSH. Downregulation of xCT may also contribute to the reduction in cellular GSH synthesis. Since GSH is also an antioxidant that can scavenge ROS [71,72], the increased GSH consumption may result in decreased resistance to oxidative stress in the liver of the *Aldh2*2* KI mice. Additionally, several reactive aldehydes including acrolein and 4-HNE have been reported to induce oxidative stress and lipid peroxidation resulting in further generation of reactive aldehydes [17,33,73,74]. Therefore, acrolein overload in the liver of *Aldh2*2* KI mice may induce oxidative stress leading to generation of lipid peroxidation-derived aldehydes including MDA and 4-HNE. The *Aldh2*2* KI mice are prone to accumulate these aldehydes since MDA and 4-HNE are also detoxified by ALDH2 [3]. This hypothesis is supported by the finding that plasma acrolein and MDA are correlated negatively with the hepatic GSH content and positively with the GSSG/GSH ratio in our model. Such a chain of reactive aldehyde accumulation due to decreased detoxification capability is considered to cause “aldehyde storm”, a vicious cycle of systemic exposure to reactive aldehydes in the *Aldh2*2* KI mice. It has been reported that reactive aldehydes form adducts with proteins and DNA, resulting in cellular damage [20,75]. Our results showed that the deposition of aldehyde adducts is present within the earliest histological lesions of the hepatocellular injury in the *Aldh2*2* KI mice after AA exposure, suggesting the involvement of the aldehyde storm in the exacerbation of acute hepatocellular injury. Furthermore, aldehyde-DNA adducts are also a key contributor to carcinogenicity [76]. The increased DNA damage in the hepatocytes of the *Aldh2*2* KI mice may imply an increased risk for hepatic cancer in the *ALDH2*2* carriers if they are chronically exposed to reactive aldehydes.

In this study, accumulation of lipid peroxidation end products MDA and 4-HNE and intracellular iron coincided or preceded with the hepatocellular injury in the *Aldh2*2* KI mice after AA exposure, suggesting that ferroptosis is involved in the exacerbation of liver injury [56,77]. The combination of GSH depletion, increased oxidative stress, and iron dysregulation is considered to induce ferroptosis in the hepatic lesion of the *Aldh2*2* KI mice. GPX4 is a central inhibitor of ferroptosis that reduces reactive polyunsaturated fatty acids phospholipid hydroperoxides to their non-reactive alcohols [56]. Thus, inactivation of GPX4 due to GSH depletion can sensitize the liver to ferroptosis even with the amount of GPX4 unchanged. Iron is incorporated into cells mainly by TFR1-mediated system and is then safely stored in ferritin nanocages or present in the form of the labile iron pool that can promote lipid peroxidation [78]. GSH is a major ligand for labile ferrous iron in the cytosol [79]and the GSH-iron complex is delivered to ferritin via poly(rC) binding protein 1 [80]. In this study, the hepatic expression of ferritin tended to decrease in the *Aldh2*2* KI mice at 120 min after AA exposure, at which iron began to accumulate in the injured hepatocytes. Decreases in iron ligand and iron storage protein may increase intracellular free iron leading to ferroptosis induction. Further investigation is required to elucidate the molecular mechanism underlying the iron dysregulation in the liver of the *Aldh2*2* KI mice after AA exposure. The protective role of ALDH2 from ferroptosis is shown in a rat model of ethanol-induced gastric ulcer [81] and is suggested in urogenital cancer [82]. To the best of our knowledge, this is the first report that suggests an involvement of ferroptosis in liver injury using an *in vivo ALDH2*2* model. Further studies using ferroptosis inhibitors ferrostatin-1 [56] and liproxstatin-1 [83] or iron chelators [84] are needed to elucidate the detailed molecular mechanisms underlying aldehyde-induced ferroptosis in our *Aldh2*2* KI mouse model. Since GSH plays a key role in oxidative stress suppression, ferroptosis suppression, and aldehyde detoxification, persistent or high consumption of tissue GSH may promote ferroptosis in *ALDH2*2* individuals, suggesting crosstalk between aldehyde metabolism and ferroptosis pathways.

Acute acrolein overload in the *Aldh2*2* KI mice resulted in tissue injury not only in the liver but also in the bone marrow, spleen, and intestines, suggesting a systemic influence of the aldehyde storm. Aldehyde storm may expose systemic organs to multiple aldehyde species. The anti-MDA clone 1F83 antibody used in this study has been reported to recognize complex aldehyde adducts consisting of MDA: MDA-acetaldehyde (M2AA-Lys)-adducts consisting of dihydropyridine derivatives [85], which have high toxicity [86] and immunogenicity [87], and MDA-formaldehyde (M2FA)-lysine adducts [87]. Our finding implies that *ALDH2*2* carriers may have a risk of systemic damage upon exposure to high levels of aldehydes exogenously and/or endogenously. Interestingly, our results showed a marked hematopoietic cell injury in the bone marrow with persistent elevation of plasma formaldehyde in the *Aldh2*2* KI mice after AA exposure. Formaldehyde has been shown to have cytotoxicity in mouse hematopoietic stem/progenitor cells in the bone marrow [52,88]. Notably, the plasma formaldehyde levels in our *Aldh2*2* KI mice after AA exposure are comparable to those reported in *Aldh2^−/−^Adh5^−/−^* mice, which exhibit perinatal lethality and lymphopenia [89], suggesting that excessive accumulation of formaldehyde alone can cause severe systemic toxicity including immunotoxicity. The elevation of plasma formaldehyde in the *Aldh2*2* KI mice is considered to be caused by a combination of ALDH2 dysfunction and ADH5 downregulation in the liver, as the two enzymes are responsible for formaldehyde detoxification [89]. Since ADH5 oxidizes S-hydroxymethylglutathione, a spontaneous adduct formed between formaldehyde and GSH [90], the detoxification of formaldehyde by ADH5 is dependent of GSH. Thus GSH depletion may lead to ADH5 dysfunction. Additionally, the bone marrow is also known for endogenous sources of formaldehyde generated through demethylation of DNA and histones and creatine metabolism pathway [91–93]. Measurement on tissue levels of multiple aldehydes from whole organs would reveal tissue susceptibility to the systemic aldehyde storm.

Interestingly, the *Aldh2*2* KI homozygous mice had higher levels of hepatic GSH and hepatic MDA than in the WT mice under physiological conditions. The abundance of GSH may be a compensatory mechanism against exposure to mild oxidative stress in the normal-appearing liver of the *Aldh2*2* KI mouse. This hypothesis is supported by the concept of “metabolic remodeling” to produce more GSH in *ALDH2*2* carriers [94,95]. Likewise, the elevation of plasma formaldehyde in the *Aldh2*2* KI mice under physiological conditions may affect certain organs such as hematopoietic system that requires formaldehyde for its function [93].

In conclusion, AA-induced acrolein overload leads to acute and systemic aldehyde storm followed by extensive ferroptosis in the *Aldh2*2* KI mice, which is probably caused by hepatic GSH depletion due to impaired aldehyde detoxification. This is the first study that reveals the crosstalk between aldehyde metabolism and ferroptosis pathways using an *in vivo* model. As humans are assumed to be routinely exposed to reactive aldehydes from the diet such as pickled vegetables and fried meat, chemicals including drugs [12] and cigarette smoke [3], and some pathophysiological conditions such as oxidative stress, the influence of reactive aldehydes on human health would be broader than expected. Further investigation is required on the health impacts after chronic exposure to reactive aldehydes in *ALDH2*2* carriers, particularly on their involvement in carcinogenesis as reported in esophageal [96] and gastric [97] cancer.

## Abbreviations

ALDH2: aldehyde dehydrogenase 2
WT: wild type
MDA: malondialdehyde
4-HNE: 4-hydroxy-trans-2-nonenal
AA: allyl alcohol
GSH: glutathione
ROS: reactive oxygen species
GPX4: glutathione peroxidase 4
KI: knock-in
PLP: periodate-lysine-paraformaldehyde
γ-H2AX: phosphorylated-histone H2AX
HMGB1: high mobility group box protein 1
TUNEL: terminal deoxynucleotidyl transferase dUTP nick end labeling
HPLC: high performance liquid chromatography
SDS: Sodium dodecyl sulfate
ADH5: alcohol dehydrogenase 5
GCLC: glutamate-cysteine ligase catalytic subunit
xCT: cystine transporter solute carrier family 7 member 11/xCT
TFR1: transferrin receptor 1
GSSG: glutathione disulfide
TBARS: thiobarbituric acid reactive substances
ALT: alanine aminotransferase

## Author contributions

**Yuki Takami:** Writing – original draft, Methodology, Investigation, Formal analysis, Data curation. **Jun Nakamura:** Conceptualization, Data curation, Funding acquisition, Investigation, Methodology, Supervision, Writing - review & editing. **Jun Katahira:** Methodology, Resources, Writing - review & editing. **Yuhei Maeda:** Methodology, Resources. **Miyuu Tanaka:** Methodology, Writing - review & editing. **Mitsuru Kuwamura:** Methodology, Supervision, Writing - review & editing. **Toshiya Okada:** Methodology, Resources, Supervision. **Takeshi Izawa:** Conceptualization, Formal analysis, Funding acquisition, Methodology, Project administration, Supervision, Writing - original draft.

---

**Supplemental Figure S1.**
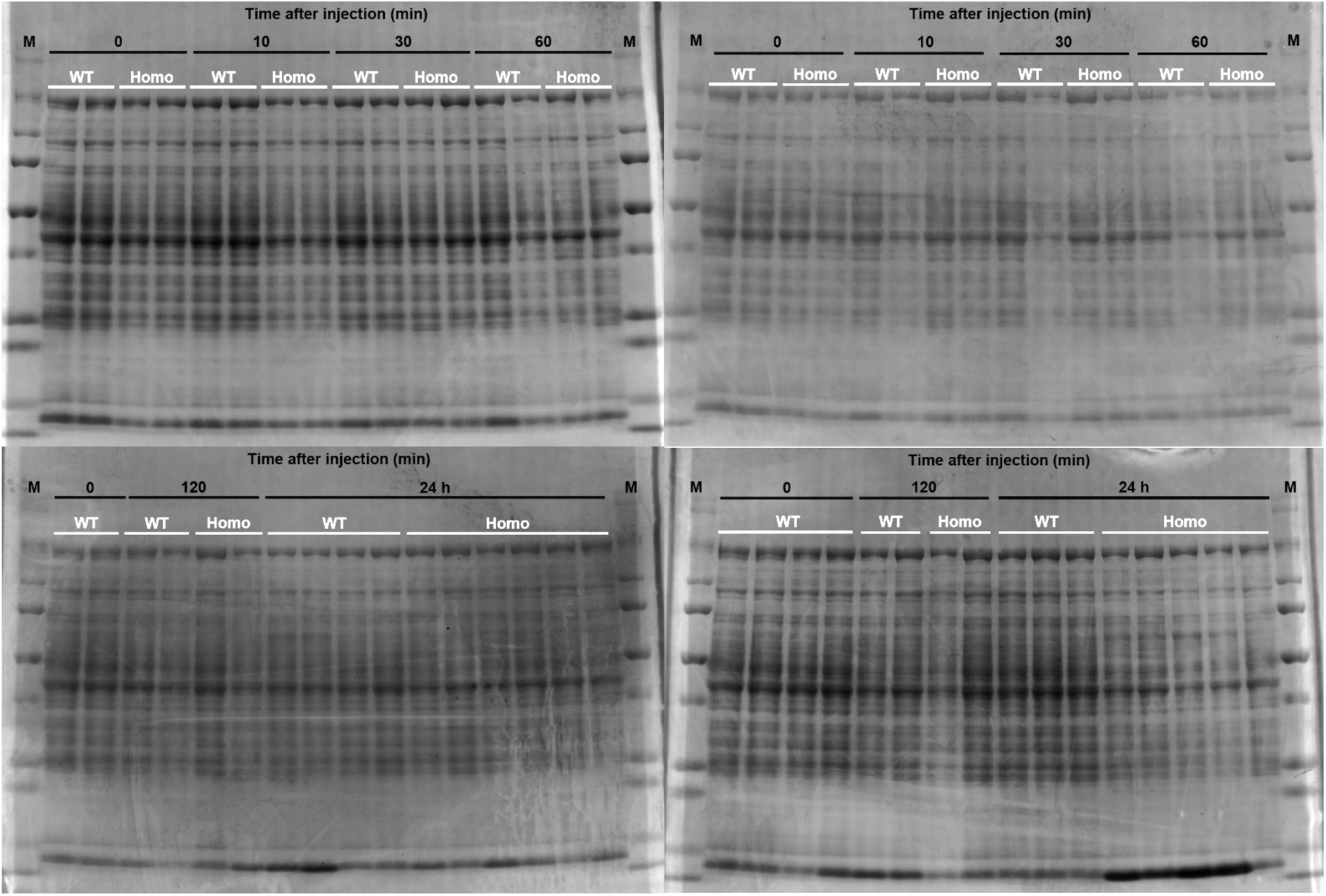
Images of the acrylamide gels stained with Coomassie brilliant blue for total protein normalization in the Western bot analysis. M, marker (#1610373, Bio-Rad, Hercules, CA, USA); WT, wild type.

**Supplemental Figure S2.**
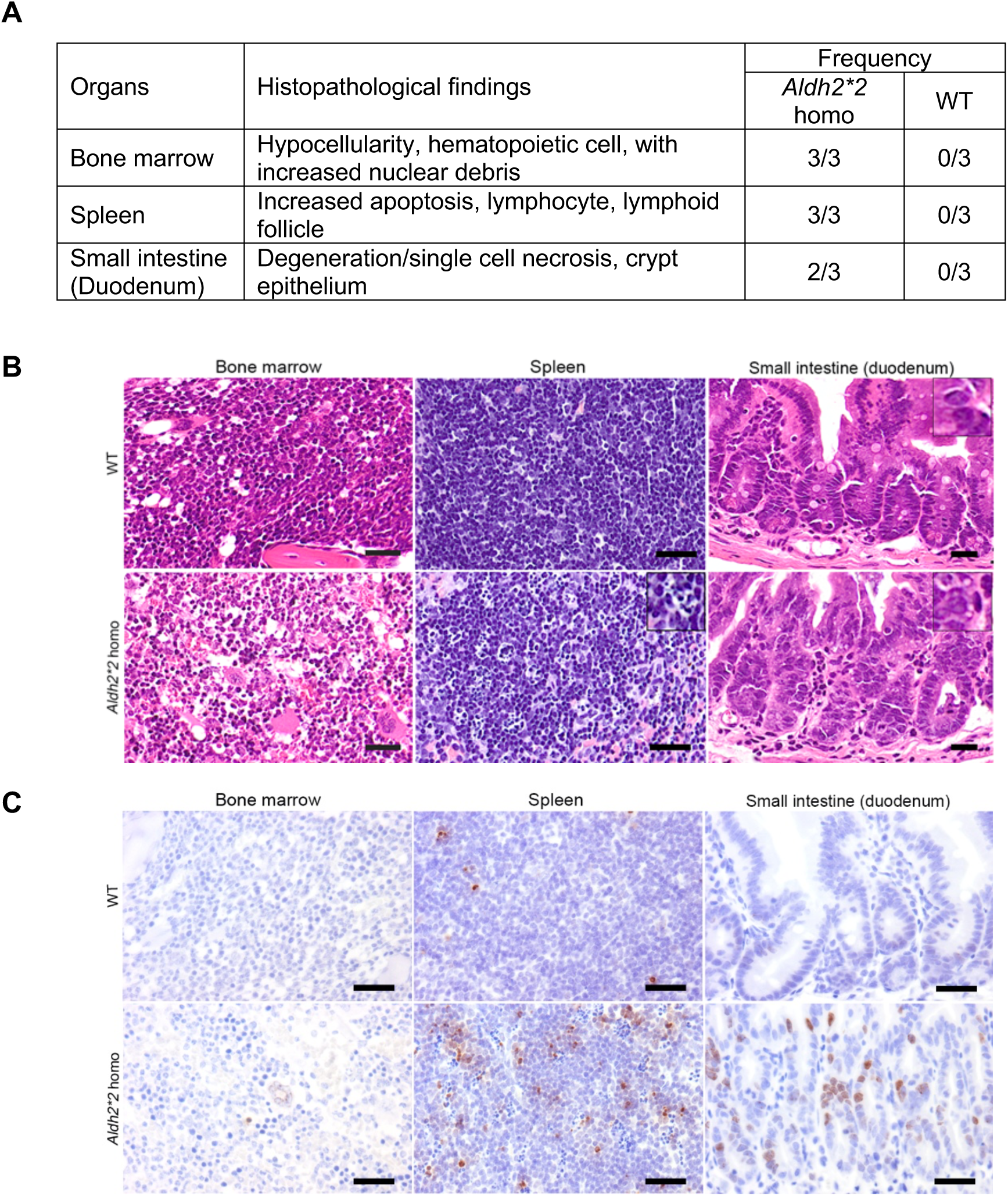
Systemic organ toxicity after allyl alcohol (AA)-induced systemic aldehyde storm in the *Aldh2*2* knock-in (KI) homozygous mice. (A) Histological findings and their frequency of bone marrow, spleen, and small intestine (duodenum) of *Aldh2*2* KI homozygous and wild-type (WT) mice at 24 h after exposure to AA at 75 mg/kg. (B) Representative images of hematoxylin and eosin stain. In the homozygous mice, the density of hematopoietic cells decreases with increased nuclear debris in the bone marrow, suggestive of acute hematopoietic cell injury. In the spleen, apoptosis of lymphocytes (inset) is markedly increased in the homozygous mice compared with that in the WT mice. The crypt epithelium of the duodenum shows vacuolar degeneration with irregular nuclear shape and scattered single cell necrosis in the homozygous mice while the crypt epithelium of WT mice appears to be normal. Bar=20 µm. (C) Immunohistochemistry for Ser139-phosphorylated histone H2AX shows an increased DNA damage in the histological lesions. Bar=30 µm.

**Supplemental Figure S3.**
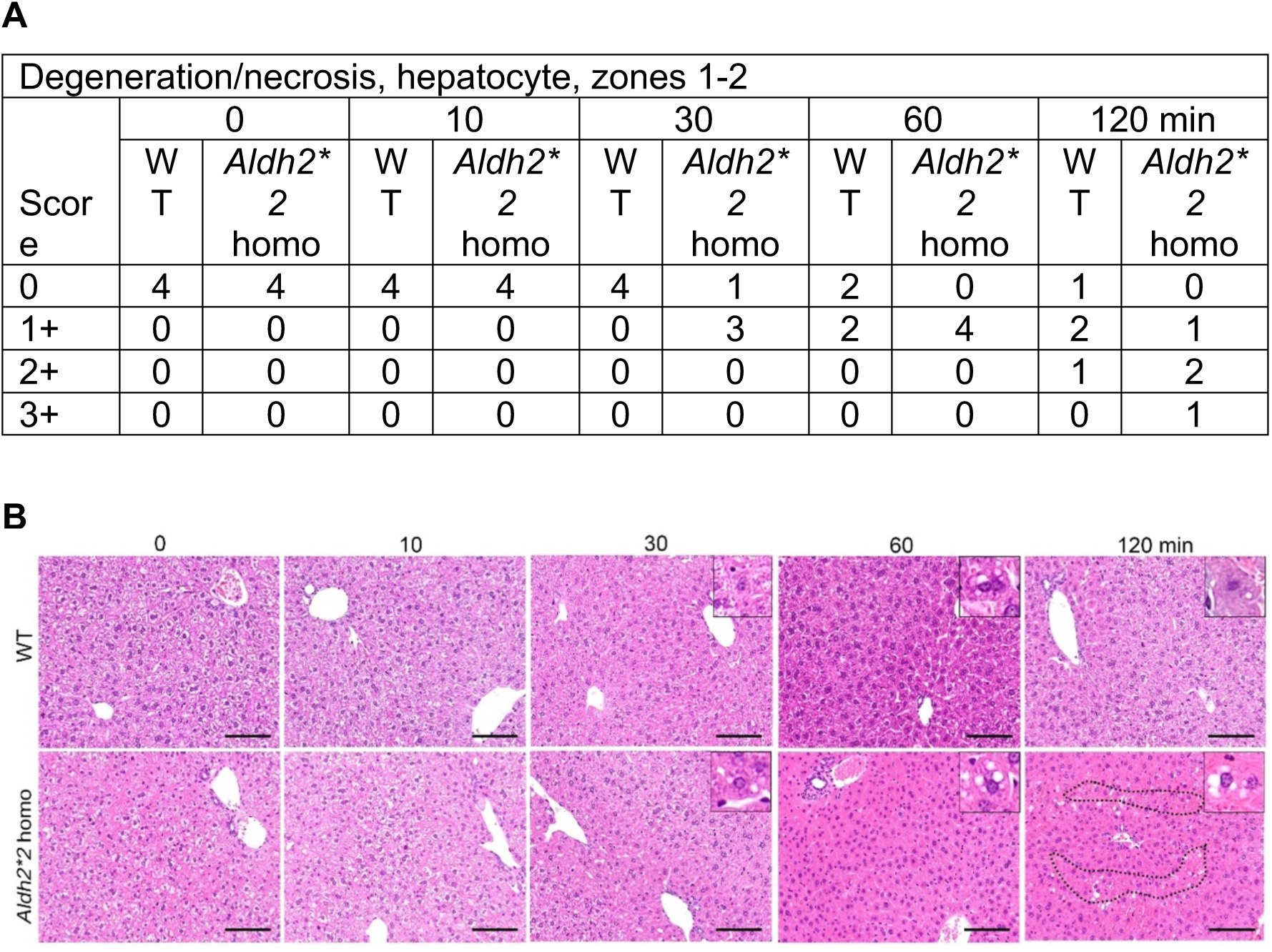
Earliest histopathological changes of the liver are detectable at 30 min after AA exposure in the *Aldh2*2* KI homozygous mice. (A) Histological score for hepatocellular injury in the hyperacute phase of the systemic aldehyde storm is described in the table. Histopathological changes of hepatocytes were graded on the scale of 0 to 3: 0 (absent), normal histological architecture; 1 (mild), focal degeneration of hepatocytes with cytoplasmic vacuoles; 2 (moderate), extensive degeneration of hepatocytes with occasional necrosis or focal coagulative necrosis of hepatocytes; 3 (severe), massive or extensive coagulative necrosis and degeneration of hepatocytes. The value represents the number of mice classified into the corresponding score. (B) Representative images of the hepatic histopathology in the WT and *Aldh2*2* KI homozygous mice after AA exposure at 75 mg/kg. Focal vacuolar degeneration of hepatocytes (insets) is observed at 30 min in the homozygous but not in the WT mice. Hepatic necrosis (encircled by dotted line) is observed at 120 min in the homozygous mice. Hematoxylin and eosin stain. Bar=100 µm.

**Supplemental Figure S4.**
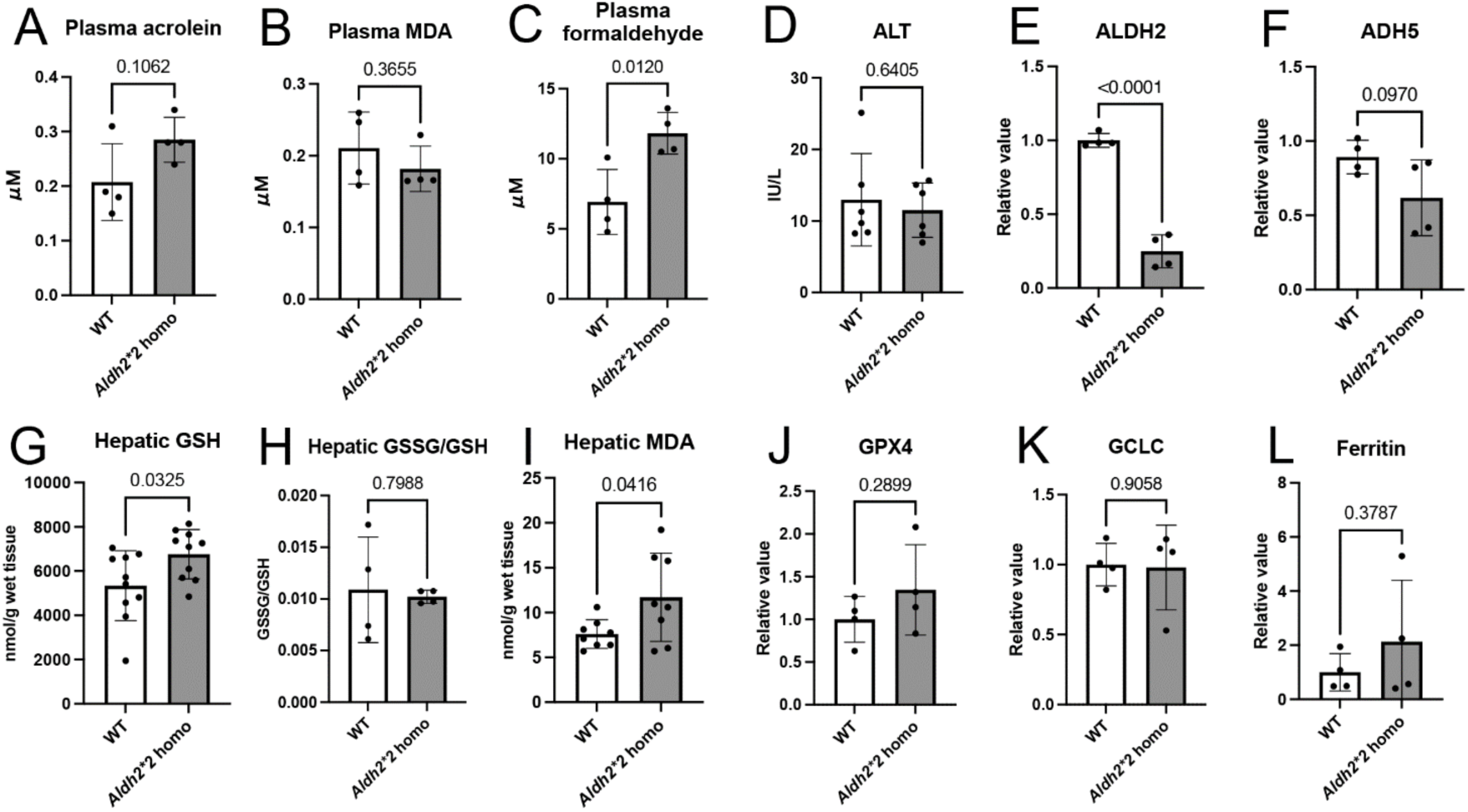
Comparison of basal physiological conditions of the liver and plasma between the WT and *Aldh2*2* KI homozygous mice. Plasma levels of reactive aldehydes (A) acrolein and (B) malondialdehyde (MDA) do not change significantly between the WT and homozygous mice, while plasma levels of (C) formaldehyde are higher in the homozygous mice than in the WT mice. (D) Serum alanine aminotransferase (ALT) does not change between the WT and homozygous mice. Hepatic expression of (E) aldehyde dehydrogenase 2 (ALDH2) decreases significantly in the homozygous mice compared with that in the WT mice. Hepatic expression of (F) alcohol dehydrogenase 5 (ADH5), a detoxification enzyme for formaldehyde, tends to decrease in the homozygous mice. Hepatic content of (G) glutathione (GSH) is greater in the homozygous than in the WT mice, with no significant difference in the (H) ratio of glutathione disulfide (GSSG) to GSH. (I) Hepatic MDA content is increased in the homozygous mice. Hepatic expression of (I) glutathione peroxidase 4 (GPX4), (K) glutamate-cysteine ligase catalytic subunit (GCLC), or (L) ferritin do not change significantly between the WT and homozygous mice. Data were analyzed by unpaired t-test by a Prism software (ver. 10.3.0; GraphPad, San Diego, CA, USA).

